# TAO Kinase 3 Overexpression Is a Poor Prognosis Marker for Lung Adenocarcinoma and Predicts Paclitaxel Resistance

**DOI:** 10.1101/622621

**Authors:** Tsung-Ching Lai, Yi-Chieh Yang, Yu-Chan Chang, Jean Chiou, Ming-Shyan Huang, Chih-Jen Yang, Peter Mu-Hsin Chang, Michael Hsiao

## Abstract

High expression of TAOK3 has been observed in several cancers, but very little about TAOK3-involved mechanisms has been reported. In this study, we investigated the role of TAOK3 in the therapeutic effect of paclitaxel treatment. Protein expression of TAOK3 was evaluated among 121 patients by using immunohistochemical staining. TAOK3 expression was estimated by analyzing the gene’s expression and its correlation with the IC50 of paclitaxel in cells. The expression of TAOK3 in lung cancer was highly correlated with poor overall survival, progression-free survival, and recurrence rate in lung cancer patients. The expression level of TAOK3 was positively correlated with paclitaxel resistance in the lung cancer cell lines. The depletion of TAOK3 could enhance the sensitivity of paclitaxel in lung cancer and vice versa. TAOK3 expression is associated with paclitaxel-resistance and could be a clinical predictor for disease recurrence and a potential therapeutic target in lung cancer.

## Introduction

Lung cancer is one of the most common cancer types worldwide and has a high mortality rate. Approximately 85% of lung cancer patients have nonsmall cell lung cancer (NSCLC) (1). For adenocarcinoma, the dominant histological subtype of NSCLC, which has increased understanding of molecular pathogenesis, has led to the success of targeted therapy. In contrast, squamous cell carcinoma tends to have different mechanisms of tumorigenesis and progression. Although surgery and chemotherapy efficiently control the disease progression, patients who have advanced NSCLC usually experience local or distant metastasis and can only be treated with chemotherapy. Many patients experience tumor recurrence and metastasis after being treated with surgery and chemotherapy. Accumulating evidence has indicated that metastasis and drug resistance are the major reasons for patient deaths.

TAOK3 is a serine/threonine-protein kinase related to the *S*. *cerevisiae* STE20-like family that acts upstream of MAPK modules. TAOK3 shares similarities with the GCK-like subfamily of STE20 kinases and can inhibit JNK activation in response to EGF (2). Downregulation of JNK signaling can speed lung tumor formation and reduce p53 protein stability in a KRas^G12D^ mouse model (3). TAOK3 has also been found to be involved in the p38 stress-activated MAPK cascade in response to DNA damage (4). Hyperphosphorylation of p38 was found in human lung cancer versus normal tissue (5, 6) and enhanced breast cancer metastasis to the lung (7, 8). Although TAOK3 is involved in both JNK and p38 signaling, the complete role of TAOK3 in cancer has not been investigated much in the past few decades. In addition, whether TAOK3 could be as a cancer biomarker is worth investigating. Furthermore, there seems to be some inconsistency in TAOK3 expression and its effect on cancer progression among different human cancer types. In prostate cancer, TAOK3 is overexpressed after androgen therapy (9), while TAOK3 is downregulated during colon cancer progression (10). To improve the prognostication of lung cancer patients, we presented the clinical and pathological roles of TAOK3 and provided a novel therapeutic strategy for lung cancer. Higher expression of TAOK3 may increase the resistance of lung cancer cells against paclitaxel. We suggest that NSCLC patients who need adjuvant chemotherapy after surgery should be tested for tumor TAOK3 expression to guide the selection of chemotherapeutic agents.

## Material and Methods

### Cell lines and media

The A549, CL1-0, CL1-5, H441, H661, H1299, PC13 and PC14 cell lines were cultured in Roswell Park Memorial Institute (RPMI) 1640 medium (Gibco, USA) supplemented with 10 mM L-glutamine and 10% fetal bovine serum. Cells were maintained in an incubator at 37°C with 5% CO_2_. Except for CL1-0 and CL1-5, which were kindly provided by Cheng-Wen Wu, the other cell lines were purchased from the American Type Culture Collection (ATCC).

### TAOK3 manipulation in cancer cell lines

TAOK3 full length cDNA was purchased from Origene. We used the gateway system to transfer TAOK3 into a lentiviral-based expression vector, plenti-6.3-bsd (Invitrogen, USA). In the TAOK3 shRNA knockdown group, we used pGRIPZ-shTAOK3-puro to reduce TAOK3 expression (Open Biosystems, USA).

### Cell viability assay

Two thousand cells were plated in 96-well plates one day before drug treatment in 100 mL medium. Cells were treated with variable concentrations by adding 1/10 volume of a ten times concentrated drug for 72 hr. Cell viability was determined by the AlamarBlue (Invitrogen, USA) assay. Fluorescent signals were recorded by an ELISA reader (PerkinElmer, USA).

### Case selection

There were a total 121 of lung cancer patients treated at the Kaohsiung Medical University Hospital of Taiwan from 1991 to 2007 in this study, consisting of 84 adenocarcinomas, 28 squamous carcinomas and 9 large cell carcinomas. All treatments for patients followed the hospital guidelines, which are annually modified by lung cancer committees in the cancer center according to the NCCN guideline and Taiwan FDA rules. All samples were all primary tumors, and no recurrent or treated lung tumors were enrolled.

Chemotherapy was applied to patients in advanced stages. Overall survival (OS) was defined as the entire period after treatment completion to death. Progression-free survival (PFS) was defined as the interval between after treatment completion and distant metastasis/recurrence/death. The study was approved by the Institutional Review Boards and the ethics committees of the involved institution (KMUHIRB-E(1) 20160099).

### TMA immunohistochemistry

Tissue microarrays were constructed with 1-mm-diameter cores from the selected formalin-fixed paraffin embedded (FFPE) tissues. Immunohistochemistry (IHC) staining was performed on 5-μm-thick sections using a Robustric Immunostainer (Ventana Medical Systems, USA). Slides were first dewaxed by heat and xylene and rehydrated in graded ethanol, followed by 30 mins antigen retrieval in pH 8.0 TRIS-EDTA buffer. The TAOK3 antibody was a polyclonal rabbit anti-human antibody (ProteinTech, USA) and was used at 1:100 for staining. The slides were subsequently counterstained by hematoxylin. The IHC staining assessment was independently observed by two pathologists blinded to the clinical information. The immunoreactivity intensity was recorded. The details of the immunostaining interpretation were described in a previous study (11).

### Analyzing TAOK3 expression and the IC50 of anti-cancer drugs

The mRNA expression levels of TAOK3 in the lung cancer cell lines were obtained by the Cancer Cell Line Encyclopedia (CCLE). The IC50 of the anti-cancer drugs were provided by the CancerDR (http://crdd.osdd.net/raghava/cancerdr/).

### Statistical analysis

All statistical calculations were performed in Excel, and the results are shown as the means ± standard deviations. Pearson’s chi-square test using a clinicopathologic table was calculated with an online tool, Chi-Square Calculator in Social Science Statistics. The Kaplan-Meier survival and Cox proportional regression hazards analysis were performed using R software. Statistical significance was accepted as a *p* value less than 0.05.

## Results

### Clinicopathological data

In our cohort, there was a total of 121 patients, including 84 lung adenocarcinomas (69.4%), 28 lung squamous carcinomas (23.1%) and 9 lung large cell carcinomas (7.4%). The patients’ age ranged from 3 to 82 years (mean age was 60.2). Approximately 50.6% were male. There were 47 patients (38.8%) who had a smoking history. Fifty-two patients (43.0%) had early stage (I–II) tumors, and 69 patients (57%) had late stage (III–IV) tumors. Seventy-seven patients (63.6%) had lymph node involvement and 35 patients had distant metastases. According to their follow-up records, 74 patients (61.2%) developed tumor recurrences or distant metastasis after treatment. There were 92 patients that died, and their average OS time was 27.4 months.

### High expression of TAOK3 was correlated with a poor prognosis and recurrence

The samples were separated into low (score 0 and 1, n=79) and high (score 2 and 3, n=42) expression groups based on TAOK3 staining intensity (Fig. 1A, B, C, and D). The correlation between the expression of TAOK3 and pathological features, including age, sex, stage, TNM stage, recurrence, chemotherapy and smoking status, was evaluated by Pearson chi-square correlation (Table 1). High expression of TAOK3 was significantly positively correlated with stage (*p*=0.0005), N stage (*p*=0.0002) and recurrence (*p*=0.0134). In survival analysis, high expression of TAOK3 was correlated with a poor prognosis for both OS (*p*=0.002) and PFS (*p*=0.001, Fig. 1E and F). In univariate Cox regression survival analysis, we found higher TAOK3 expression increased the risk of an event 1.89 times relative to low expression (*p*=0.003) for OS and PFS. In the two major types of NSCLC, adenocarcinoma and squamous cell carcinoma, high expression of TAOK3 was significantly correlated with a poor prognosis for both OS (P=0.019) and PFS (*p*=0.032), especially for adenocarcinoma (Fig. S1, Table S1 and S2). In addition, we also observed similar results for the correlation of TAOK3 high expression and poor prognostic outcome for lung adenocarcinoma patients from the online database, Kaplan-Meier Plotter (Fig. S2) (12).

**Table 1.**
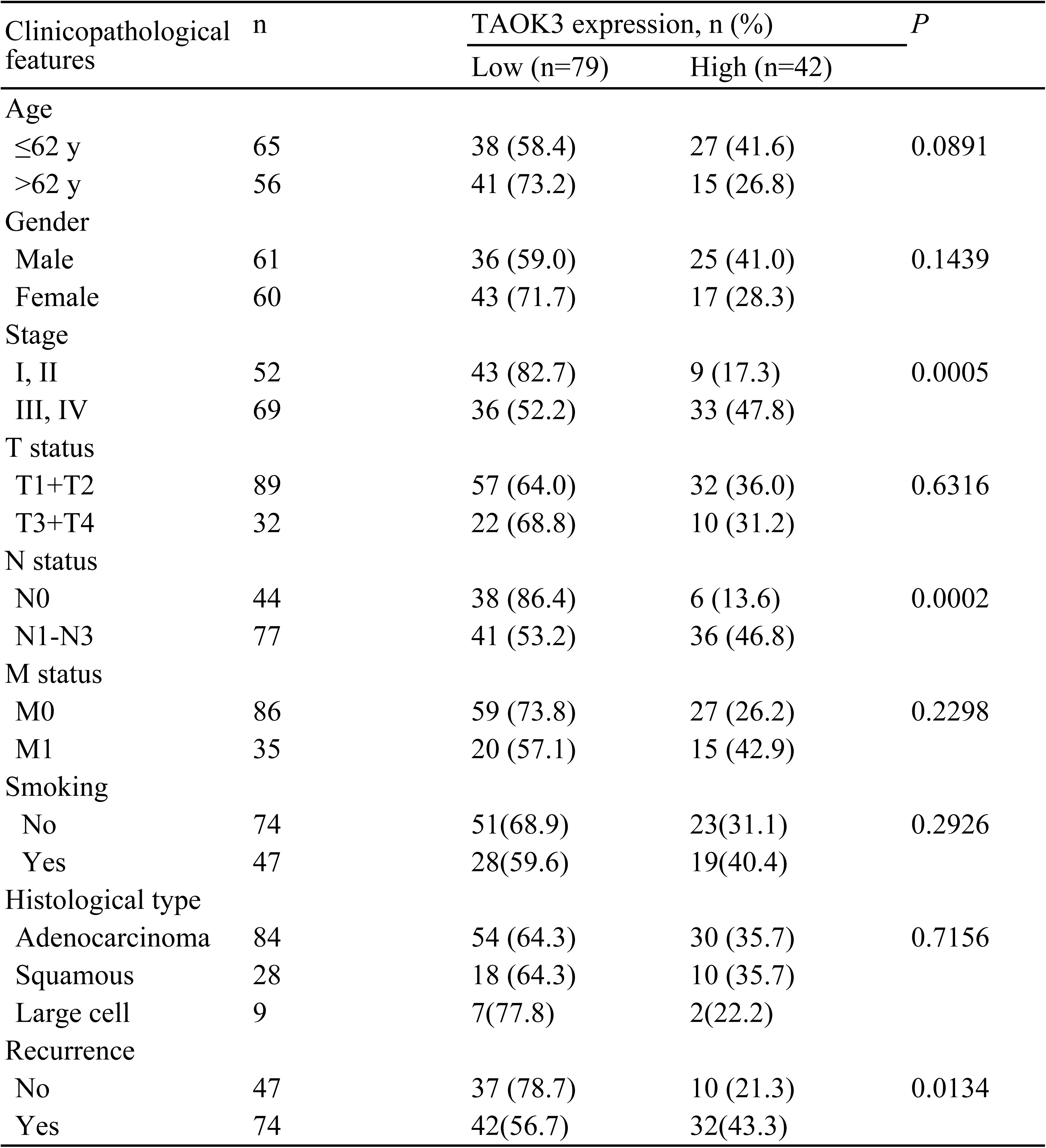
Clinicopathological features of TAOK3 expression in lung cancer patients

**Figure 1.**
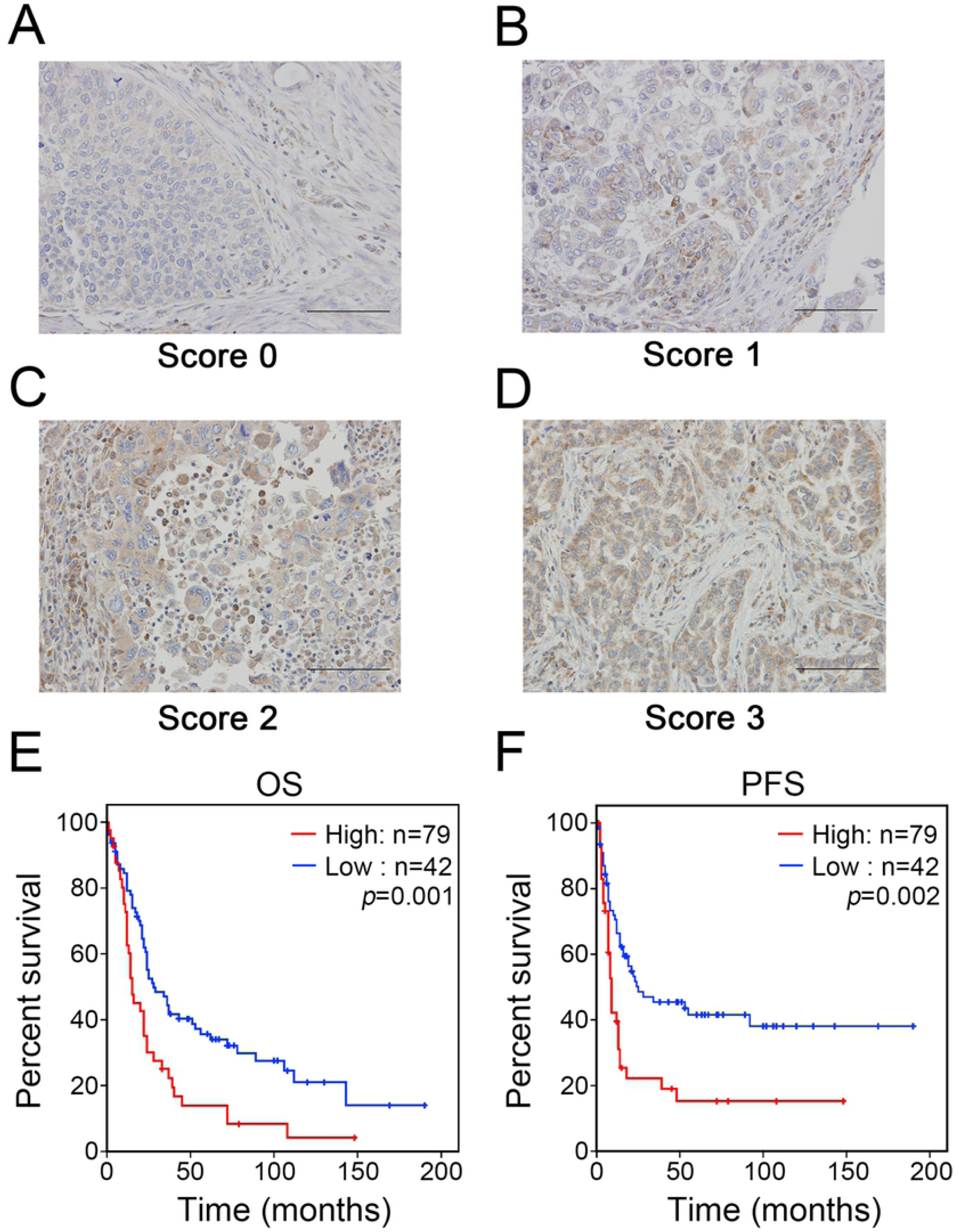
TAOK3 expression pattern in lung cancer patients. (A) The scoring pattern from 0 to 3 represents the level of TAOK3 expression from no, weak, medium and strong in lung cancer cells. (B) Kaplan-Meier curves of overall survival and progression-free survival for TAOK3 expression status in NSCLC patients.

We further evaluated the hazard ratio changes with Cox regression survival analysis in our lung cancer cohort to determine the prognostic significance among all pathologic factors. In both univariate Cox survival analyses of OS and PFS, we found that T stage, N stage, M stage and TAOK3 expression were all significantly correlated with a higher risk of death of lung cancer patients (Table 2). In multivariate Cox survival analysis, N staging showed a significant correlation with both PFS and OS. In addition, M staging was still an important factor for OS (Fig. 2 and Table 2). However, the correlation between TAOK3 expression and death was weaker in both multivariate Cox regression analyses.

**Table 2.** Cox regression survival analysis Progress free survival (PFS) - Univariate

| Factor | Hazard ratio | 95% CI | P value |
| --- | --- | --- | --- |
| T staging | 2.381 | 1.443-3.928 | 0.001 |
| N staging | 3.737 | 2.093-6.673 | <0.001 |
| M staging | 2.314 | 1.422-3.766 | 0.001 |
| TAOK3 | 2.133 | 1.335-3.408 | 0.002 |

Progress free survival (PFS) - Multivariate
| Factor | Hazard ratio | 95% CI | P value |
| --- | --- | --- | --- |
| T staging | 1.554 | 0.917-2.634 | 0.101 |
| N staging | 2.815 | 1.515-5.230 | 0.001 |
| M staging | 1.451 | 0.861-2.448 | 0.162 |
| TAOK3 | 1.456 | 0.891-2.379 | 0.134 |

Overall survival (OS) - Univariate
| Factor | Hazard ratio | 95% CI | P value |
| --- | --- | --- | --- |
| T staging | 2.519 | 1.601-3.963 | <0.001 |
| N staging | 2.868 | 1.787-4.603 | <0.001 |
| M staging | 2.768 | 1.776-4.315 | <0.001 |
| TAOK3 | 1.891 | 1.241-2.882 | 0.003 |

Overall survival (OS) - Multivariate
| Factor | Hazard ratio | 95% CI | P value |
| --- | --- | --- | --- |
| T staging | 1.589 | 0.978-2.581 | 0.061 |
| N staging | 2.108 | 1.263-3.520 | 0.004 |
| M staging | 1.765 | 1.079-2.886 | 0.024 |
| TAOK3 | 1.272 | 0.809-2.000 | 0.297 |

**Figure 2.**
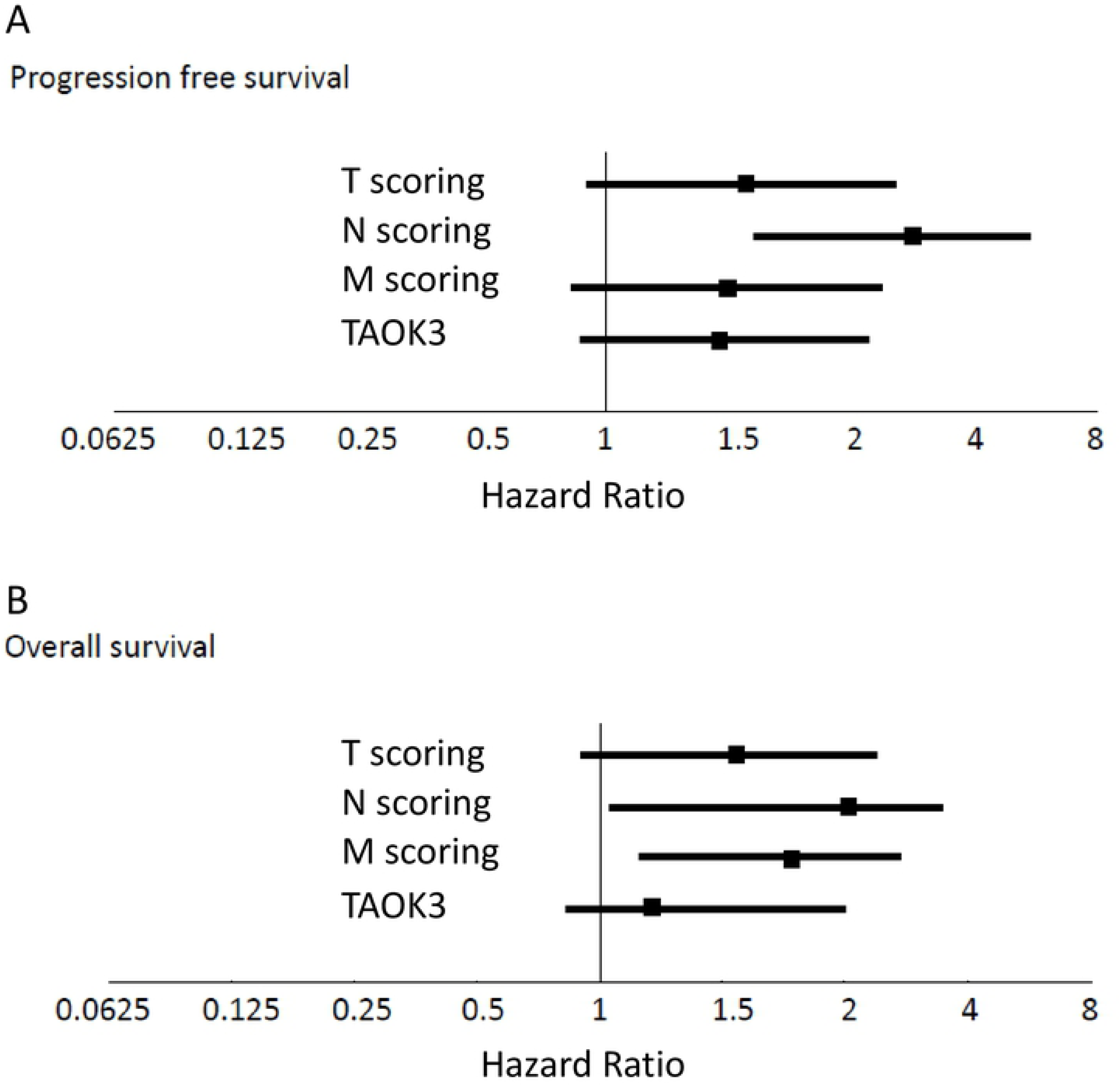
Forest plots present the hazard ratios of (A) progress-free survival and (B) overall survival from multivariate Cox regression survival analysis.

### The expression of TAOK3 was positively correlated with paclitaxel resistance in lung cancer cell lines

Currently, there are three major chemotherapeutic drugs that are combined with cisplatin for lung cancer adjuvant treatment: gemcitabine, paclitaxel and vinorelbine. By analyzing drug responses by levels of TAOK3 expression, we found there was a significant correlation with paclitaxel sensitivity (*r*=0.724, *p*=0.0179) in 10 human lung cancer cell lines but not with gemcitabine or vinorelbine (Fig. 3A and Fig. S3). Interestingly, among the human lung cancer cell lines, high endogenously expressing TAOK3 cell lines (H441 and H1299) also exhibited paclitaxel resistance (Fig. 3B). While overexpression of TAOK3 increased paclitaxel resistance, knockdown of TAOK3 sensitized lung cancer cells to paclitaxel by approximately 6 times in H1299 and 4 times in H441 (Fig. 3C). On the other hand, it upregulated TAOK3 in the low endogenous TAOK3 cell lines (PC13 and PC14) enhanced paclitaxel resistance by approximately 4 times.

**Figure 3.**
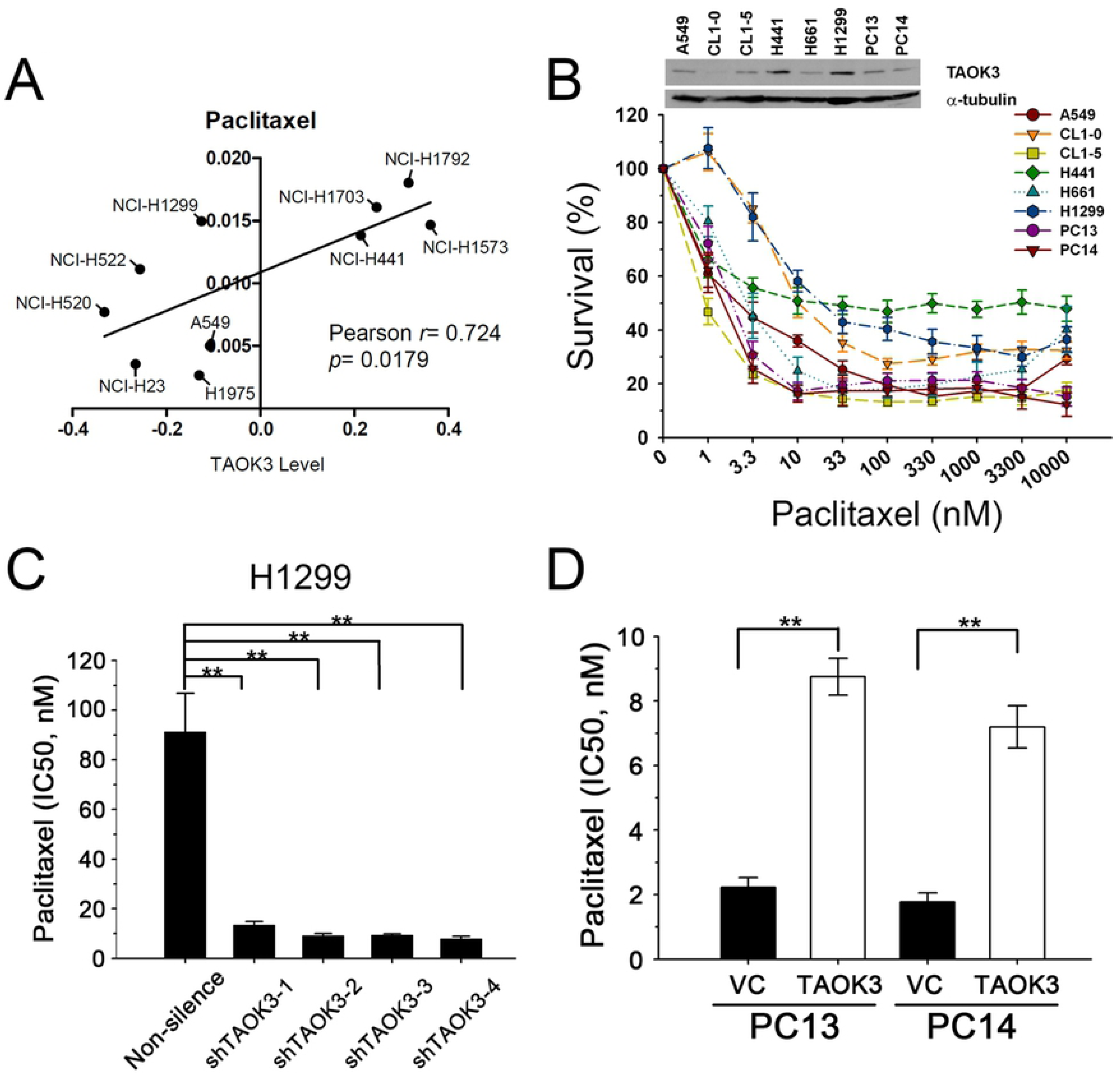
The expression level of TAOK3 in cells correlates with paclitaxel sensitivity. (A and B) Analyzing the correlation of TAOK3 mRNA (A) and protein (B) expression level and the IC50 doses of paclitaxel in lung cancer cell lines. (C) Paclitaxel sensitivity of H1299 cells with TAOK3 depletion. The results were compared to the nonsilencing sequence control. (D) Paclitaxel sensitivity of PC13 and PC14 with TAOK3 overexpressed. The results were compared to the empty vector control. \**p*<0.05, \*\**p*<0.01

## Discussion

In this study, we first showed that overexpression of TAOK3 was correlated with advanced N status and a poor prognosis. *In vitro* study also revealed that overexpression of TAOK3 increased resistance against paclitaxel. Since traditional chemotherapy is still the standard adjuvant therapy for NSCLC and paclitaxel is one of the most common drugs used in combination with platinum, this study result provided us with not only a prognostic marker but also a predictive marker for the choice of platinum-based regimens.

Currently, the most important prognostic factor for NSCLC is the “Stage” by the American Joint Committee on Cancer (AJCC) TMN staging system obtained by combining the status of the primary tumor (T), local lymph node involvement (N), and distant metastasis (M) to determine the final stage status (13, 14). In the era of multimodality team approaches for the treatment of NSCLC, the most critical issue is to identify potentially occult lymph node (LN) involvement, which will change early T1N0M0 stage IA disease to at least stage IIB, and this change will influence not only the prognosis but also the choice of whether adjuvant therapy will be given. To determine the N status more precisely, the use of comprehensive or selective mediastinal LN dissection is still controversial (15). PET/CT scanning is an alternative choice, but its resolution and sensitivity will decrease for smaller LNs (14). Furthermore, in a recent study of PET/CT negative T1N0M0 NSCLC patients after lobectomy, occult N metastases were detected in 9.6% (43 of 449) of the patients (16). This study also suggested that higher glucose uptake by the primary tumor (PET-SUVmax) may indicate more LN involvement (16). Other clinical pathologic factors have been proposed, such as a primary tumor larger than 2 cm, and one study mentioned that adenocarcinoma may be more likely to have LN involvement than squamous cell carcinoma (17); however, another study found the opposite (16). In our study, we identified that TAOK3 is specifically correlated with N status instead of T or M.

Few studies have evaluated TAOK3 in regard to the prognosis of NSCLC. TAOK3 was reported to inhibit the tumor suppressive function of JNK, and it induces p38 signaling (18). These two alterations of MAPK pathways were reported to enhance lung nodule formation in allotransplantation tumors of a transgenic mouse model (3, 8). In our clinicopathological analysis, overexpression of TAOK3 was most strongly correlated with two metastatic indications, N status and recurrence (Table 1). This result also gives us a hint that for early stage NSCLC, TAOK3 expression level in the primary tumor may serve as a biomarker for PET/CT scanning or prompt further mediastinal LN dissection.

Adjuvant C/T is recommended for stage II–IIIA NSCLC patients since both OS and PFS benefits have been observed (19, 20). Traditional platinum-based chemotherapy is still the standard regimen, and there is no survival difference among different combinations of navelbine, taxane, tegafur/uracil, or other drugs. Although EGFR-TKIs have been proven to prolong PFS in metastatic NSCLC patients harboring EGFR mutations, their benefits during adjuvant therapy have not been well established (20, 21). In addition, there are currently no predictive biomarkers for selecting adjuvant regimens since *KRAS*, p53, ERCC1, β-tubulin and others have all been analyzed in large clinical trials, but no significant differences in treatment outcome have been observed (22-25). In this study, we accidentally discovered that TAOK3 expression is specifically associated with paclitaxel resistance. Since we could not evaluate the correlation between TAOK3 expression and outcomes of patients treated with adjuvant paclitaxel from the current cohort, its role as a novel predictive biomarker for adjuvant taxane-based chemotherapy regimens needs to be further validated. Therefore, due to the predictable role of TAOK3 in the taxane therapeutic effect on lung cancer patients, the level of TAOK3 in lung cancer patients is worth checking before selecting the appropriate chemotherapy drug. A pilot population of patients would need to be enrolled and undergo fine-needle aspiration to test their tumor TAOK3 level and their follow-up treatment response. These tests would apply our basic bench findings to the clinical setting.

Taxane resistance is well known to be due to pumping it out of cancer cells by ATP-binding cassette (ABC) transporters, including p-glycoprotein and multidrug-resistant proteins expression (26, 27). The relationship between TAOK3 expression and taxane resistance has not been previously evaluated. The possible mechanism of TAOK3 induced paclitaxel resistance may be correlated with its role in stress responses by regulating p38/MAPK pathways. It has been reported that depletion of TAOK could impair damage-induced G2/M arrest (4). On the other hand, Taxol induced G2/M arrest resulting in cell apoptosis in the past decade (28). Collectively, it is reasonable that we speculate about whether TAOK3 could help cells to survive under taxane treatment.

## Conclusions

In conclusion, this study identified a new biomarker, TAOK3, for use as both a prognostic factor and predictive marker for paclitaxel. It is especially helpful for locally advanced NSCLC patients who could have TAOK3 IHC staining performed on resected tumors and these results may act as a reference to help physicians make decisions about further adjuvant chemotherapy regimens. Larger clinical studies are warranted to validate its clinical use in the future.

### Abbreviations

TAOK3: serine/threonine-protein kinase TAO3
NSCLC: nonsmall cell lung cancer
FFPE: formalin-fixed paraffin embedding (FFPE)
IHC: immunohistochemistry
NCCN: National Comprehensive Cancer Network
JNK: c-Jun N-terminal kinase
CCLE: Cancer Cell Line Encyclopedia
PET/CT: positron emission tomography/computed tomography

## Declarations

## Acknowledgements

We would like to thank Ms. Tracy Tsai for her assistance with the immunohistochemistry.

## Availability of data and materials

The detailed information about the data generated or analyzed in this study is included in this published article and some patient data is included in the supplementary file.

## Authors’ contributions

TCL and YCC conceived and designed the study; TCL, YCC, JC, and YCY performed the experiments and collected data; MSH, JC, and CJY were responsible for the analysis and interpretation of data; TCL, YCC, YCY, and MH drafted the manuscript; and CJY, PMC, and MH supervised the study and revised the manuscript. All authors read and approved the final manuscript.

## Consent for publication

Not applicable.

## Ethics approval and consent to participate

Tumor samples were harvested with informed consent from patients who were undergoing surgical resection at the Department of Surgery of Kaohsiung Medical Hospital, The Kaohsiung Medical University in Taiwan. The study was approved by Institutional Review Board and written consent was obtained from patients prior to their inclusion.

